# Phased Mutations and Complex Rearrangements in Human Prostate Cancer Cell Lines through Linked-Read Whole Genome Sequencing

**DOI:** 10.1101/2021.08.09.455584

**Authors:** Minh-Tam Pham, Harshath Gupta, Anuj Gupta, Ajay Vaghasia, Alyza Skaist, McKinzie A. Garrison, Jonathan B. Coulter, Michael C. Haffner, William B. Isaacs, Sarah J. Wheelan, William G. Nelson, Srinivasan Yegnasubramanian

## Abstract

A limited number of cell lines have fueled the majority of preclinical Prostate cancer (PCa) research. Despite tremendous effort in characterizing their molecular profiles, comprehensive whole genome sequencing with allelic phasing of somatic genome alterations has not been undertaken to date. Here, we utilized whole genome Linked-read sequencing to obtain haplotype information from the seven most commonly used PCa cell lines (PC3, LNCaP, DU145, CWR22Rv1, VCaP, LAPC4, MDA-PCa-2b), four castrate resistant (CR) subclones (LNCaP_Abl, LNCaP_C42b, VCaP-CR, LAPC4-CR), and an immortalized prostate epithelial line RWPE-1. Phasing of mutations allowed derivation of “Gene-level Haplotype” to assess whether a gene harbored heterozygous mutations in one or both alleles, providing a comprehensive catalogue of mono or bi-allelically inactivated genes. Phased structural variant analysis allowed identification of complex rearrangement chains consistent with chromothripsis and chromoplexy, with breakpoints occurred across a single allele, providing further evidence that complex SVs occurred in a concerted event, rather than through accumulation of multiple independent rearrangements. Additionally, comparison of parental and CR subclones revealed previously known and novel genomic alterations associated with the CR clones. This study therefore comprehensively characterized phased genomic alterations in the commonly used PCa cell lines and provided a useful resource for future cancer research.

## INTRODUCTION

Cancer cell line models are vital resources in cancer research. For the PCa field, derivation of cultured cell lines from human prostatic cancers has proven very challenging, with fewer than 20 unique parental cell lines established in the past half-century of research (1, 2, 3). These cell lines have become precious resources for preclinical studies of PCa; a simple query of PubMed revealed >23,000 publications making use of these cell lines. Interestingly, the top 7 most used of these cell lines (PC3, LNCaP, DU145, CWR22Rv1, VCaP, LAPC4, MDA-PCa-2b) accounted for >99% of all of these citations Given how valuable these cell line models are, many efforts have been made to characterize the genomic features and molecular profiles of these cell lines. For example, in two landmark studies, Bokhoven et al. and Beheshti et al. performed spectral karyotyping and molecular profiling and revealed evidence of both chromosomal instability and gene mutations as oncogenic drivers(4, 5). These earlier studies shed light on the mutational burden and oncogenic mutational and structural drivers of classic PCa cell lines.

Advances in next generation sequencing (NGS) have fueled large scale genomic studies using whole exome and/or whole genome sequencing of >1,000 PCa genomes, identifying driver genes recurrently mutated in human PCas (6, 7, 8). These studies have allowed identification of recurrently mutated PCa driver genes even when they are relatively infrequent across individuals (7). Prostate cancer genomic studies have also allowed identification of driver structural alterations in PCa (9, 10, 11, 12). Baca et al analyzed the anatomy of structural variants (SV) in PCa and coined the term chromoplexy to describe complex SVs that likely occur through a transcription-coupled process (12). Quigley et al. established the relationship between driver mutations and SV patterns in metastatic PCa (12). Together, a better picture of the mutational profile of PCa is coming to light, informing novel therapeutic and diagnostic strategies for the deadly disease. Next generation sequencing has also allowed ever more comprehensive and deep characterizations of the ways in which the PCa cell lines resemble or differ from primary human cancer tissues. One recent study even carried out 70X whole genome sequencing of two commonly used PCa cell lines LNCaP and PC3 with de novo assembly (13). However, it is still unclear to what extent the common PCa cell lines recapitulate the genomic mutational and structural alterations found in patient samples.

Recent development of long read sequencing and linked read sequencing has allowed relatively comprehensive assessment of haplotypes and even end-to-end assembly of human genomes (11, 14, 15, 16). The long-range genomic information enables haplotyping of alleles to phase blocks up to tens of mega base pairs (Mbp) long (15, 16). Combining the power of sequencing fidelity of traditional Illumina seq and emulsion based barcoding technologies to generate libraries of small reads from high molecular weight DNA enabling assembly of the larger DNA segments, linked-read sequencing has enabled phasing of variants, and greater power for determination of structural alterations. Such linked read sequencing has not yet been used to determine the genomic alterations in the commonly used PCa cell lines.

Here we report a comprehensive analysis of phased mutations and SVs determined through linked-read sequencing of high molecular weight DNA from the 7 most commonly used PCa cell lines. This analysis allowed assessment of mono-versus bi-allelic gene mutations and phasing of complex structural variant breakpoints. Additionally, juxtaposition between the parental and castrate resistant (CR) subclones for 3 of the cell lines revealed SVs and mutations potentially associated with CR progression. Comparison of the genomic alterations in these cell lines to those identified in large scale PCa genome sequencing studies revealed that this series of cell lines harbored a significant fraction of the recurrent putative driver mutations, and complex structural rearrangement patterns observed in human PCa. Therefore, this study serves as a comprehensive compilation of genomic features for the most commonly studied PCa cell lines, representing a valuable resource for the field.

## MATERIALS AND METHODS

### Cell lines and culturing methods

Cell lines were obtained and cultured in media as instructed by ATCC with the exception of LAPC4, LAPC4-CR, and VCaP-CR. LAPC4 was cultured in IMDM with 10% fetal bovine serum and 1nM R1881, and LAPC4-CR and VCaP-CR were cultured in 10% charcoal striped fetal bovine serum (Thermo Fisher A3160401) RPMI with added 1X B27 supplement (Thermo Fisher 17504044). Frozen pellets were used for mycoplasma testing and STR genotyping (GenePrint 10 System) performed by the Johns Hopkins Genetic Resource Core Facility. STR percent match was calculated using the second Master vs query algorithm (17) based on ATCC query STR data. With cell lines that do not have ATCC query STR data, the STR profile of the parental cell line was used for reference. All cell lines were mycoplasma free and passed the a priori defined STR genotype threshold of 70%.

### High Molecular weight (HMW) DNA Extraction

HMW DNA was extracted following the “Salting out” demonstrated protocol described by 10X Genomics (Pleasanton, CA) Technical Support. Briefly, two million cells were pelleted and lysed overnight at 37°C in Lysis buffer containing 2mM EDTA, 300mM NaCl, 8mM Tris-HCl, 125ug/mL Proteinase K, and .5% SDS. HMW DNA was precipitated using 1M NaCl, washed in absolute ethanol, and resuspended overnight at 4 □ C in TE buffer. Quality control for HMW DNA was performed with the TapeStation 4200 system (Agilent). All samples contained more than 80% of DNA of greater than 60kb in length.

### Barcoding, library preparation, and sequencing

1 (+/-0.2) ng of DNA was quantified by Qubit Broad Range (Thermo Q32850) and used for GEM creation and barcoded library generation using the 10X Chromium Genome protocol. Resulting library fragment sizes were determined using the DNA 1000 and 2100 Kit BioAnalyzer (Agilent Technologies), and indexed with Chromium i7 Sample Index Plate (Catalog number 220103 from 10xGenomics). The index library libraries were sequenced to 30X-50X average coverage on an Illumina HiSeqX platform using paired-end 150 bp x 150 bp reads. The resulting sequencing BCL files were processed by the Long Ranger Pipeline (14) (10X Genomics) for alignment, variant discovery, and phasing.

### Analysis for phased variants

Only passed variants from phased_variants.vcf files produced by Long Ranger were annotated using the vcf2maf (18) pipeline and the Uniport consensus reference. To filter for likely somatic variants, only variants not described in the gnomAD database or those with gnomAD_AF of less than .1% were retained. To limit to variants with potential protein functional consequences, any variants annotated as “IGR” (integenic), “Silent”, “Targeted_Region” under Variant classification were filtered out. Categories that started with “regulatory_region_variant”, “TF_binding_site_variant”, “splice_donor_variant”, “splice_region_variant”, “coding_sequence_variant” under Consequence were also filtered out. For the list of genes frequently implicated in cancer, the list of 97 genes recurrently mutated in PCa as identified by Armenia et al. as well as all Cosmic Tier 1(19) 576 genes were used as a reference list of genes recurrently mutated in PCa and cancer in general. *Cosmic-Longtail* is the union of the two aforementioned lists, consisting of 629 genes. Gene-level haplotype was inferred through a custom R (version 4.0.0) script that used the haplotype (column GT) and phase set (column PS) information provided by Long Ranger in phase_variants.vcf files. In short, any gene with at least one mutation with GT=“1|1” was categorized as “Hemi/homozygous”, and any gene with only one heterozygous (GT=“0|1” or “1|0”) mutation was categorized as “Monoallelic”. For any gene with multiple heterozygous mutations, if all mutations were on the same phase set and belonged to the same haplotype (GT= “0|1” or “1|0”), it was categorized as “Multi-monoallelic”. If all mutations were on the same phase set and there was at least one mutation belonging to the opposite allele, it was categorized as “Biallelic heterozygous”. In cases where multiple heterozygous mutations did not belong to the same phase set or were unphased (GT=“0/1”), the genes were categorized as “no info”.

### Analysis for phased structural variants

Phased structural variants (SV) were defined as large SVs calls in large_sv_calls.bedpe file from Long Ranger that have phased set information (PS1/PS2). A custom R script, termed ChainLink, allowed inference of phased structural variant breakpoints as follows: SVs were chained together based on both shared phased set and shared haplotype for each breakpoint. Each SV chain contained at least 2 SVs. ChainFinder (12) was used in parallel using phased SVs from all 12 cell lines to determine background breakpoint frequency. “Copy number type” was set as “Seq” and Deletion_threshold was set at 2. Other parameters were left as default.

SnpEff 4.5.1 (RRID:SCR_005191) (20) was used to annotate the closest gene to each breakpoint coordinate.

### Graphics

All graphs were prepared using 10x Genomics Loupe Browser 4.1.0 and the R packages Rcircos (1.2.1) (RRID:SCR_003310) (21) and ggplot2 (3.3.5) (RRID:SCR_014601), assembled on Adobe Illustrator (24.1.1) (RRID:SCR_010279).

### Packages

Long Ranger 2.2 (14), tidyverse (22), Rcircos 1.2.1 (21), SnpEff 4.5.1 (20), ChainFinder (12)

## RESULTS

### Linked-read sequencing of human PCa cell lines

We performed linked read sequencing of high molecular weight DNA from 11 PCa cell lines and one immortalized non-malignant prostatic epithelial cell line RWPE-1 (Fig. 1A; Table S1). The cancer cell lines included three widely-used androgen deprivation sensitive cell lines (LAPC4, VCaP, and LNCaP), and their respective androgen deprivation insensitive subclones LAPC4-CR (23), VCaP-CR (23), LNCaP_Abl (24) and LNCaP_C42b (25). Additional androgen receptor (AR) positive cell lines included CWR22Rv1, an androgen deprivation sensitive line derived from a xenograft model established from a primary adenocarcinoma, and MDA-PCa-2b derived from a patient of African American ancestry. Remaining cell lines included two commonly used AR-negative prostatic carcinoma lines, PC3 and DU145. All cell lines were confirmed to be devoid of mycoplasma contamination and to have close genetic matching of short tandem repeat (STR) genetic markers to the previously documented STR genotyping information (Table S1). For all cell lines, >80% of the extracted DNA had high molecular weight >60 kbp as measured by TapeStation Bioanalyzer (Table S2). Next generation sequencing of linked-read libraries generated from these samples resulted in 35X to 54X average genome coverage (Table S2). Initial quality analysis of the linked-read sequencing data revealed that on average, 53 to 97 linked reads could be phased to high molecular weight DNA molecules with a calculated average size of 59 kb to 144 kb (Fig. 1B; Table S2). These data could be used to assemble haplotype phased blocks typically ranging from 10s to >100 megabase (Mb), allowing phasing of >98% of all single nucleotide polymorphisms (SNPs) and genes for all cell lines (Fig. 1B, C; Table S2).

**Figure 1.**
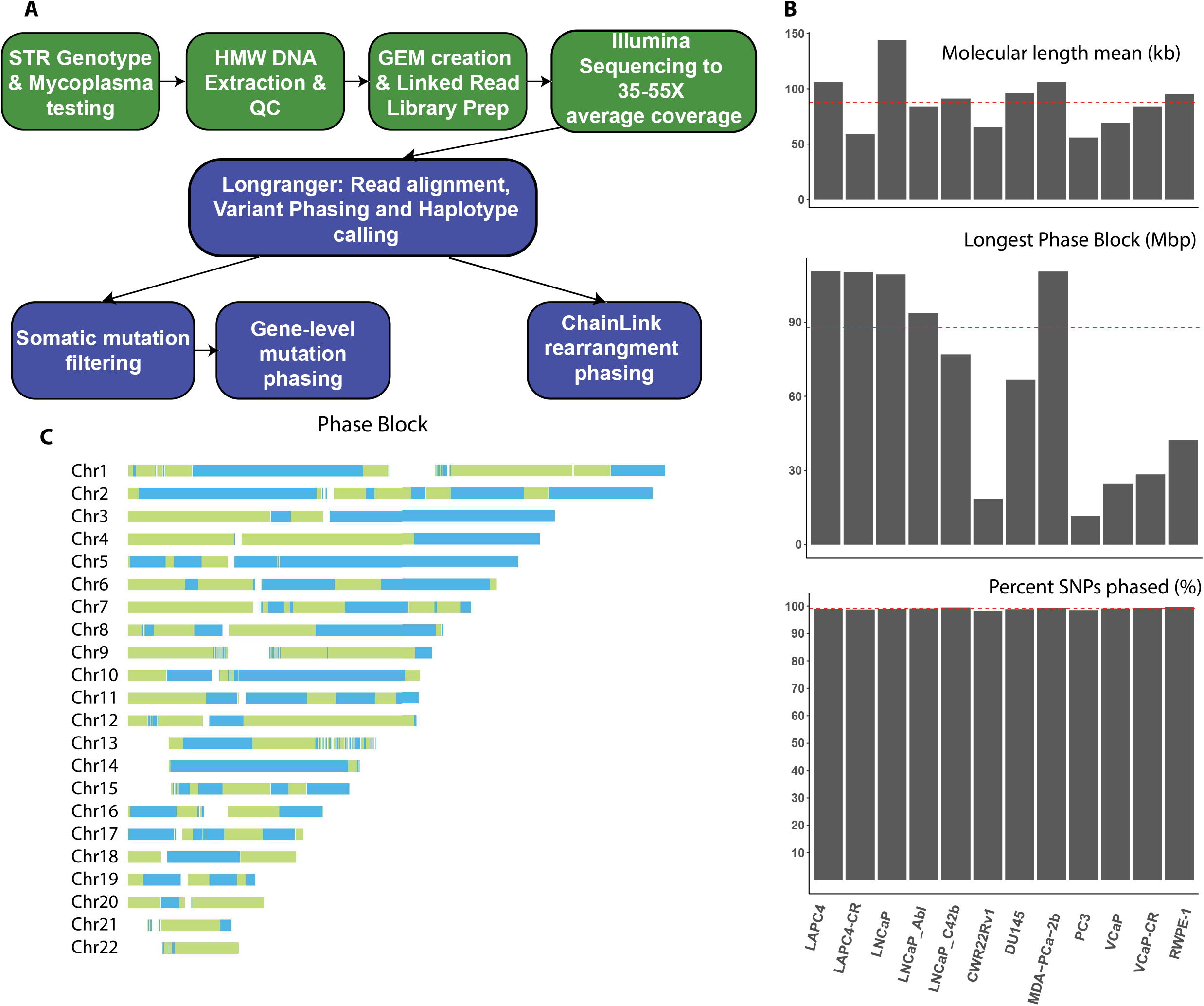
Performance of linked read sequencing of PCa cell lines. **(A)** Schematic of workflow and analysis steps. Green boxes represent wet lab experiments; blue boxes represent computational analysis. **(B)** Performance metrics of linked read sequencing for each cell line, including computed mean molecular length in kilobases (kb), length of longest phased block in mega-bases (Mb), and percent of SNPs phased. Dotted red lines represent mean values across cell lines. **(C)** Representative phase block ideogram from LAPC4 showing each phase block as a single-colored and contiguous stretch of chromosome with alternating colors representing neighboring phase blocks.

### Identification of somatic mutations in the PCa cell lines

We assessed likely somatic mutations, defined as nucleotide variants (SNV) and small insertions and deletions, by filtering for mutations that exhibited very low population allele frequencies (<0.1%) in the Genome Aggregation Database (gnomAD)(26). Somatic mutations accounted for 0.88% of all variants detected. As expected, we confirmed that cell lines with known microsatellite instability (MSI) and mismatch repair deficiency (LNCaP, LAPC4 and their CR clones, CWR22Rv1, DU145, and MDA-PCa-2b (27, 28)) exhibited high mutation rates (Fig. 2A, Table S3). We found that MSI high cell lines harbor 2500 to 5000 coding somatic mutations, with CR clones from LNCaP acquiring additional 20-30% of mutations compared to parental, mostly missense. Unsurprisingly, microsatellite stable (MSS) VCaP and PC3(27) cells exhibited lower mutational burdens, with ≤ 4 somatic mutations per Mbp genome, and less than 1000 coding mutations (Table S3, Fig. 2A). Amongst the MSI high cell lines, 70-75% of mutations were missense, with 12-25% indels. MSS cell lines had less than 70% missense mutations, with 20% or more indels, and more than 10% splice site mutations (Fig. 2A).

**Figure 2:**
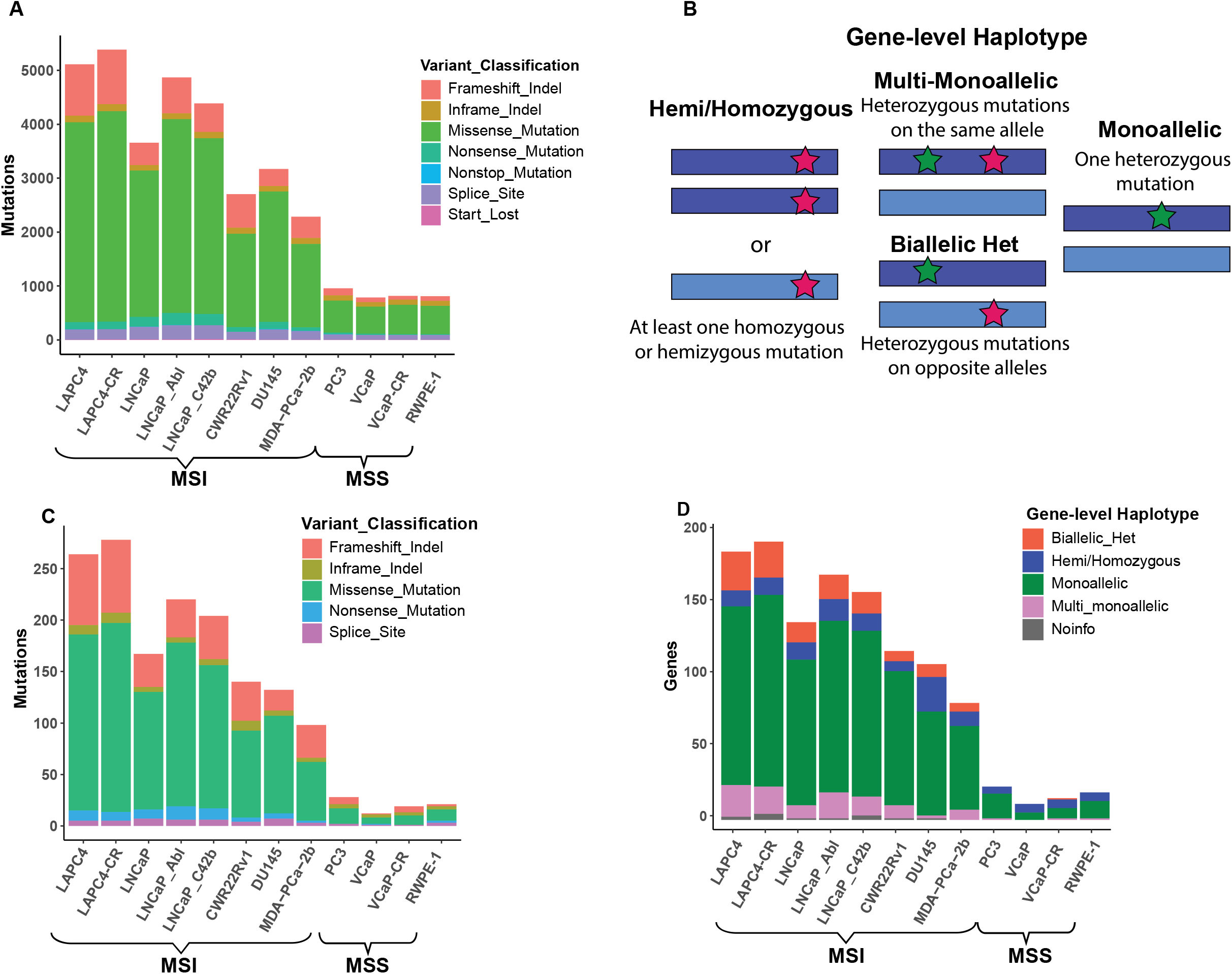
Phasing of somatic mutations to determine Gene-level Haplotype. **(A)** Somatic mutation counts of cell lines stratified by MSI status, with all mutations stratified by protein alteration classification. **(B)** Gene-level Haplotype distinguishes between genes with multiple heterozygous mutations on the same or different alleles. Dark blue or light blue bars represent each allele for a given gene. Red and green stars indicate position of a mutation. **(C)** Somatic mutations on *Cosmic-Longtail* gene list, stratified by variant classification and **(D)** Gene-level haplotype.

### Gene-level phasing of somatic mutations to distinguish between complete versus single copy gene mutations

Linked read sequencing allowed phasing of heterozygous mutations, thus revealing additional information regarding the allelic mutation status at the gene level. Using the phasing information derived from the linked read sequencing analysis, we developed the concept “Gene-level haplotype” for each gene to identify mutations in each gene at each haplotype to determine whether a given gene is affected at a single or both alleles (Fig. 2B). First, any genes with at least one homozygous or hemizygous mutation were designated as “Hemi/homozygous”. Any gene with one heterozygous mutation was called “Monoallelic”. “Multimonoallelic” describes a gene with multiple mutations on only one allele. “Biallelic heterozygous” describes a gene with at least two heterozygous mutations on both alleles, thus leaving no wild-type allele at the gene level. In cases where a gene contained heterozygous mutations that were not properly assigned to a haplotype, or the phase block ended within a gene, it would be labelled as “no info”.

Gene-level haplotypes across all genes showed that MSS cell lines had a high percentage of hemi/homozygous mutations on genes (20-40%), indicating loss of heterozygosity (LOH) events as a result of copy number or structural changes (Fig. S1A). Ten to seventeen percent of all genes mutated had multiple heterozygous mutations, which were resolved into multi-monoallelic (< 10% for all cell lines), and biallelic heterozygous (8-15% for MSI cell lines and nearly absent in MSS cell lines) with phase information. Monoallelic genes accounted for 65-75% of mutated genes in all cell lines (Fig. S1A). Leveraging prior studies and databases cataloguing recurrently mutated driver genes in human cancers (29), including PCa (7), we compiled a list of putative PCa driver genes representing the union of genes annotated in the Cosmic tier 1 list(29) and those identified in Armenia et al., (7) and called this the **Cosmic-Longtail** list. Among these, we saw a high percentage of frameshift mutations across all cell lines (Fig. 2C). MSI cell lines showed higher percentage of genes inactivated by biallelic heterozygous mutations (Fig. 2D). In contrast, amongst 629 genes in the **Cosmic-Longtail** genes, VCaP showed 50% of genes were hemi/homozygous and 50% monoallelic. The high percentage of genes mutated in hemi/homozygous manner indicates loss of heterozygosity and chromosomal instability is more likely the driving force for carcinogenesis in this cell line.

### Recurrently mutated driver genes in human PCa are highly represented in the PCa cell lines

We further examined more closely genes recurrently mutated in human PCa only as profiled by Armenia et al (7) that we called *Longtail* gene list. Out of the 97 genes in the *Longtail* list, we found 87 genes to be mutated in our panel of 12 cell lines. To make sure any missense mutations in these genes were potential cancer drivers, we filtered to include only those mutations identified as clinically relevant driver mutations as compiled by the Cancer Mutation Census (CMC) project under COSMIC (29). We further included frameshift, splice-site, and nonsense inactivating mutations within the *Longtail* list, yielding 58 genes with putative driver mutations found in our panel of cell lines (Fig. 3A).

**Figure 3:**
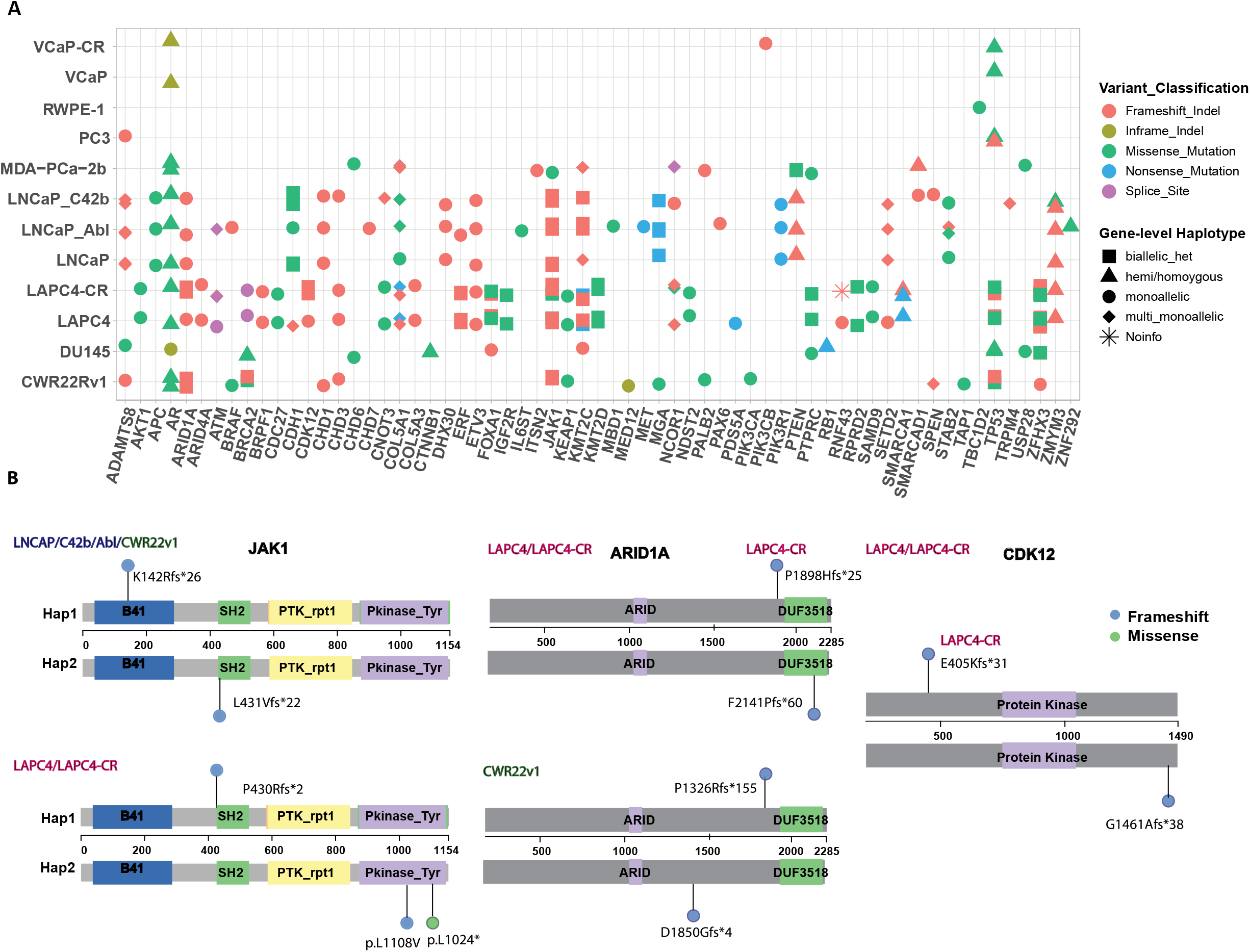
Prostate cancer cell lines harbor a large fraction of recurrent driver mutations found in human PCas. **(A)** Cell line mutations in the *Longtail* panel of cancer driver genes (7) that are recurrently mutated in human PCa, with variant classification for each mutation, and Gene-level haplotype for each gene annotated for each cell line. **(B)** Lollipop plots showing locations of *JAK1, ARID1A*, and *CDK12* biallelic heterozygous mutations relative to positions of Pfam domains. Colors of the lollipop head indicate the variant classification for each mutation.

Compared to all genes, the *Longtail* list showed a higher fraction of frameshift mutations (Fig. S1B). Gene-level haplotypes allowed identification of several biallelic heterozygous alterations, including in *CDK12, ARID1A*, and *JAK1* (Fig. 3B). Cyclin dependent kinase 12 is a transcription kinase that is responsible for phosphorylating the second serine residue of RNA Polymerase II C-terminal domain, in addition to being involved in RNA splicing and transcriptional elongation of long genes (9, 30). *CDK12* mutation denotes a distinct subclass of PCa characterized by tandem duplication and high neoantigen burden (9, 10). In our cell lines, LAPC4 and its CR clone both have G1461Afs*38 (Mutation ID COSM2837928). This confirmed somatic mutation occurred 14 times in the CMC, and is the most common frameshift *CDK12* mutation, recurrently found in colon adenocarcinoma, pancreatic ductal carcinoma, and gastric adenocarcinoma (19). It is interesting to note that this frameshift mutation led to a lengthening of the protein by 9 amino acids, along with alteration of the last 30 amino acids of the protein. Another recurrent somatic frameshift mutation in *CDK12* T1463Nfs*50 (COSM1382846) also alters the last 28 amino acids and adds an additional 22 amino acids to the protein. It is thus intriguing to speculate that these recurrent mutations may lead to altered function rather than just loss of function. LAPC4-CR lost the other copy of *CDK12* from an E405Kfs*31 mutation, which truncated the remaining two thirds of the protein.

Two other examples to note were the SWI/SNF subunit *ARID1A* and the tyrosine kinase *JAK1*. CWR22Rv1 had heterozygous mutations p.P1326Rfs*155 (COSM133029) and p.D1850Gfs*4 (COSM133004) on opposite alleles of *ARID1A*, both found a couple times in human cancer (29). LAPC4 and LAPC4-CR had p.F2141Pfs*60 (COSM51429 for p.F2141Sfs*59) and LAPC4-CR had an additional frameshift mutation on the other allele p.P1898Hfs*25 (COSM1639822) thus inactivating both alleles. Janus Kinase 1 (*JAK1*) is a tyrosine kinase implicated in immune and apoptosis evasion (31). LNCaP, its CR clones, and CWR22Rv1 all had p.L431Vfs*22 (COSM41842) and p.K142Rfs*26 (COSM1639943), both of which are recurrent *JAK1* mutations in human cancer. The exact two *JAK1* mutations on two separate alleles occurred in both CWR22Rv1 and LNCaP (and its CR sub-lines). LAPC4 and LAPC4-CR shared p.P430Rfs*2 (COSM1560531) and had additional mutations at p.L1108V and p.L1024* on the kinase domain. Loss of function mutations in *JAK1* is common among MSI cell lines and associated with reduced interferon response (32). In our limited cell lines cohort, biallelic heterozygous mutations on *JAK1* happen in many MSI cell lines, representing a stepwise tumorigenic process to slowly evade apoptosis and immune surveillance in MSI cancer cells (32).

### Structural variants disrupt many Cosmic-Longtail genes

Structural variants and CNVs also disrupted multiple genes in the *Cosmic-Longtail* list. Structural variant breakpoints landed on exons, introns, or within 5kb from 24 genes in the *Cosmic-Longtail* list (Table S7). Notably, the tumor suppressor *FHIT* was involved in multiple unique duplications and deletions in many cell lines. Specifically, in DU145, two homozygous deletions of 42 kb and 30 kb were found in introns (Table S7) and two single copy deletions (CN=1) of 260 kb and 1.26 Mb were found towards the beginning and the end of the *FHIT* gene (Table S8). LNCaP_Abl and C42b both shared a 300 kb duplication spanning from the middle of *FHIT* towards the middle of its neighbor gene *PTPRG* (Table S7). LNCaP_C42b also carried an additional 139 kb deletion near the start of the gene, and a 397 kb duplication towards the end of the gene (Table S7). In LAPC4, a 50 kb deletion was found in a middle intron of FHIT (Table S7), while LAPC4-CR harbored two duplications of 113 kb and 772 kb in the middle of the gene, accompanied by a translocation with chromosome 1 fusing the FHIT gene to the *ESRRG* gene locus (Table S7). LAPC4-CR also harbored a 6.1Mb gene deletion (CN=1) towards the beginning of *FHIT* (Table S8). These observations were consistent with prior observations that *FHIT* locus is one the most fragile in epithelial cells and it is one of the earliest and most frequently altered in human cancer (33). Additionally, the continued SV and CNV accumulation in CR clones suggests *FHIT* might play a role not only in cancer initiation but also progression.

MSS cell lines PC3 and VCaP accumulated more alterations on putative cancer driver genes through SVs and CNVs. A previously documented 840 kb deletion on chromosome 10 resulted in total *PTEN* loss in PC3 (Table S9) (13), a 50 kb deletion on chromosome 19 fused exon 2 of the MAPK regulator and tumor suppressor **CIC** with exon 14 of *TMEM145* (Table S7), and a heterozygous 55 kb deletion on chromosome 6 removed the first 2 exons of the type I human leukocyte antigen *HLA-A* (Table S7). PC3 also carried an LOH 133 kb duplication on chromosome 14 which duplicated the 5th exon of the DNA damage repair gene *RAD51B* (Table S7). In VCaP, beside the known *TMPRSS2-ERG* fusion, an inverse translocation on chromosome 5 fused the first 3 exons of the oncogenic transcription factor *EBF1* to the latter half of the pseudogene *ADGRV1*, potentially introducing a fusion gene (Table S7). We also observed a 460 kb deep deletion (CN=0) on chromosome 8 causing total deletion of the tumor suppressor phosphatase *PPP2R2A* in VCaP, and a full 620kb deep deletion (CN=0) on chromosome 4 that completely deleted a series of genes involved in chemokine signaling and inflammatory pathways *CXCL3, CXCL2, MTHFD2L*, and *AREG* (Table S9).

### Phasing of structural alterations in PCa cell lines

Prostate cancers have been associated with having numerous structural variants, including highly recurrent rearrangements involving *ETS* transcription factors, often part of complex rearrangement patterns such as chromoplexy or chromothripsis (11, 12, 34, 35, 36). The rearrangements in chromoplexy and chromothripsis are thought to arise in a coordinated manner, as opposed to sequential formation of independently arising rearrangements (12, 34). Chromoplexy involves multiple intra- and inter-chromosomal rearrangement breakpoints that likely form in a single event (12, 35). Chromothripsis describes a phenomenon of a large number of rearrangements occurring within a chromosome often associated with alternating chromosomal gains and deletions, thought to occur in a single catastrophic rearrangement event (10, 36).

We next examined structural variants in the PCa cell lines using the Linked-read sequencing data. Linked read sequencing has been shown to have greater power to detect large SVs than conventional paired-end short read sequencing, since several reads with shared molecular barcodes can contribute to the identification of a rearrangement present within the original long DNA molecule. By combining rearrangement breakpoint information with the associated haplotype information of each segment of the breakpoint, we reasoned that it should also be possible to infer the phasing of multiple breakpoints to the same allele to identify complex rearrangements consistent with chromoplexy or chromothripsis. We developed a pipeline called ChainLink to infer clustered SVs by combining phase information and SV identification (Fig. 4A). A “chain” was defined as at least 2 SVs having at least one breakpoint each sharing both the same phased set and the same haplotype. SVs with breakpoints belonging to 2 different phase sets bridged different phase sets into the same chain.

**Figure 4:**
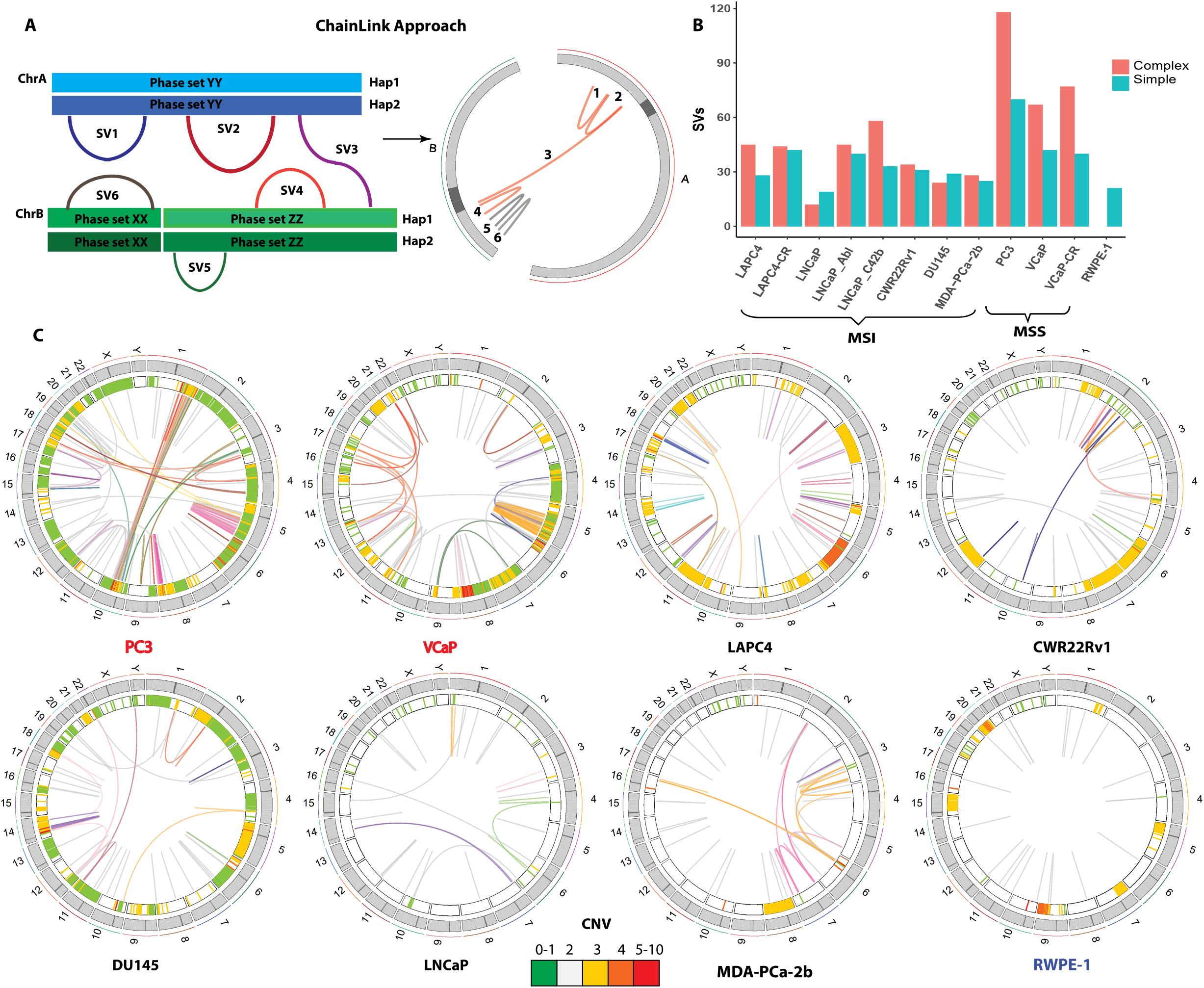
Identification and phasing of structural alterations: **(A)** The Chainlink approach uses phase information for each breakpoint to chain together complex SVs. In this schematic, SV 1, 2, 3, 4 on two different chromosomes are chained together into a complex SV cluster based on phase information on each breakpoint; SV 5 and 6 are single SVs. **(B)** Number of SVs classified as simple or complex in each cell line. **(C)** Circos plots representing CNV and SVs on the 8 parental cell lines in this study: MSI stable cell lines PC3 and VCaP (red), MSI high cell lines DU145, LNCaP, LAPC4, CWR22Rv1, MDA-PCa-2b (black) and non-malignant immortalized prostatic epithelium RWPE-1 (navy). Heatmap track beneath chromosome ideograms represents CNV with colors representing copy number. Innermost link track represents large SVs: complex SVs are colored by chain; simple SVs are gray.

Using this method, we identified complex rearrangements in all PCa cell lines, ranging from 3 chains in LNCaP to 11 chains in VCaP-CR (Fig. 4B, C). The highest number of SVs in one chain was 34, and modal number of SVs per chain was 3 (Fig. S3C). The non-cancer line RWPE-1 notably did not have any complex SVs (Fig. 4B, C). Many of these complex rearrangements were consistent with chromoplexy (present in several cell lines), and chromothripsis (most notable in PC3 and VCaP). Intriguingly, in contrast with the high number of mutations (Fig. 2A), MSI cell lines exhibit less SVs compared to MS stable cell lines (Fig. 4B,C; Fig. S2A,B).

We compared the performance of the ChainLink method to a previous method called ChainFinder (12) (Fig. S3). ChainFinder uses rearrangement breakpoints identified by conventional paired-end sequencing data to infer chained rearrangements based on comparing breakpoint locations to an expected distribution of breakpoints if they had arisen independently. Using ChainFinder with the same sequencing data, but ignoring the molecular barcode information, we found that ChainFinder identified fewer chains, and fewer structural variants per chain, relative to ChainLink. Additionally, ChainLink identified all chains called by ChainFinder (Fig. S3A, B, C). These analyses suggested that the use of phasing information from linked-read sequencing can increase the sensitivity of identifying rearrangements within complex chained rearrangements compared to more indirect statistical methods that do not incorporate such phasing information.

Using ChainLink, we were able to decipher the complete genomic anatomy of the complex SV producing the *TMPRSS2-ERG* fusion gene in VCaP cells (Fig. 5A,B; Table S5). It was previously known that the 3Mbp sequence between the *TMPRSS2* and *ERG* genes was rearranged to other genomic segments in this cell line, rather than being deleted (37); however, the precise genomic anatomy of rearrangements involving this 3Mbp stretch was not well understood. Here we show that the∼3Mbs between *TMPRSS2* and *ERG* was broken up into two parts, with one rearranged with sequences on chromosomes 16 and 17, and the other rearranged with sequences on chromosomes 16 and 12. All together the rearrangement involving this ∼3 Mb segment harbored 6 structural variants in a highly complex genomic anatomy (Fig. 5A,B). Parallel ChainFinder analysis only called three of these structural variants and could not fully decipher the complex anatomy of the rearrangements associated with this fusion gene, likely because of the large distance (1Mbp-2Mbp) between the breakpoints.

**Figure 5:**
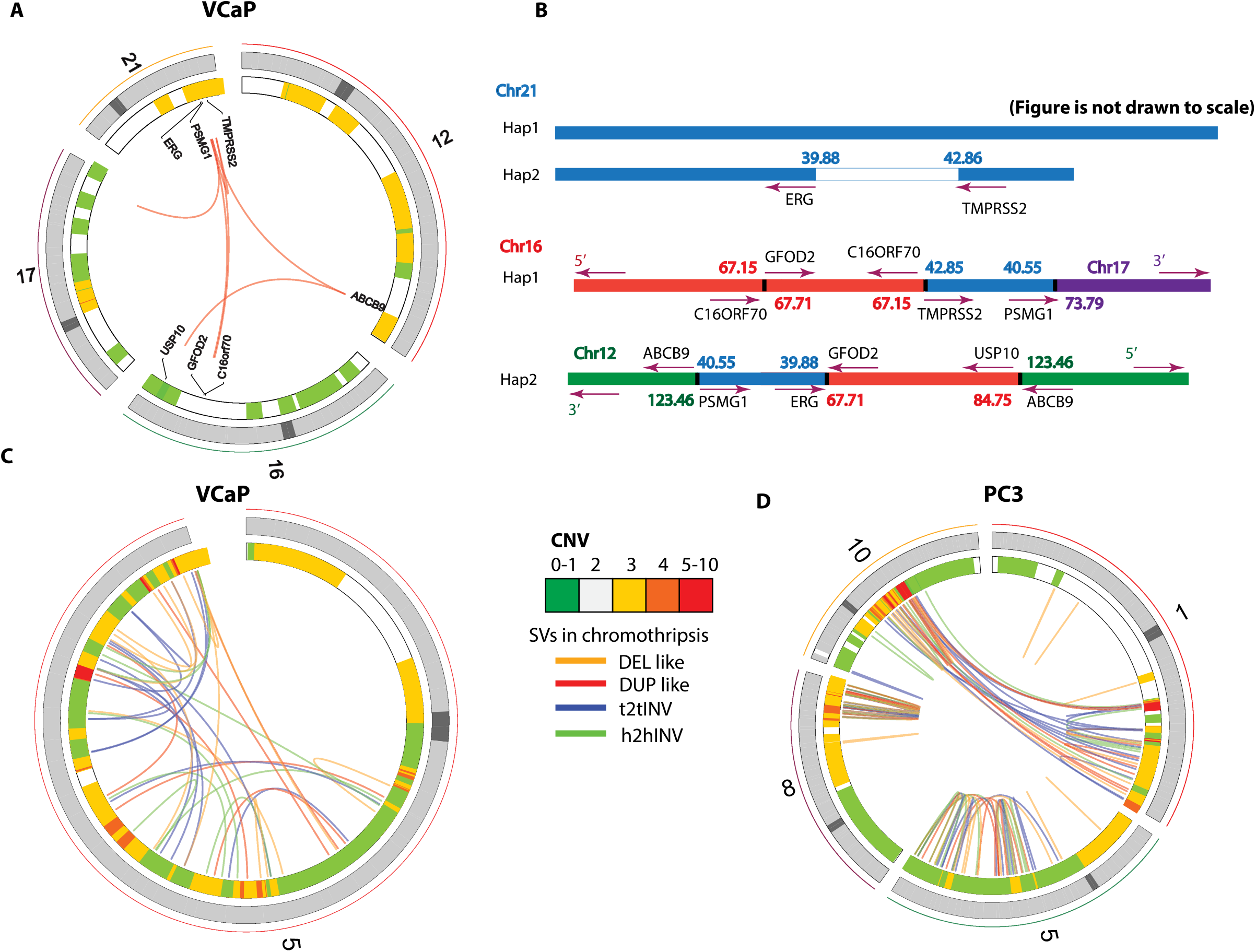
Highlights of complex chromoplexy and chromothripsis rearrangements in VCaP and PC3. (A) Circos plot of the chromoplexy event associated with the *TMPRSS2-ERG* fusion gene formation in VCaP. Genes interrupted by these SVs are labeled on the innermost track. (B) Genomic anatomy of the breakpoints involved in the chromoplexy event leading to the *TMPRSS2-ERG* fusion gene in VCaP and detailed rearrangement configuration of the 3Mb sequence between the *TMPRSS2-ERG* genomic breakpoint. Genomic coordinates are in Mbp per reference genome hg19. **(C)** Rearrangements and copy number alterations involved in a chromothripsis event on chromosome 5 in VCaP, and **(D)**, on chromosomes 5, 8, 1, and 10 in PC3. Heatmap tracks beneath chromosome bands represent CNV with colors representing copy number. Innermost link track joins the genomic breakpoints of large SVs. **(C**,**D)** Color of SV links indicates orientation of breakpoints as deletion like (DEL like), duplication like (DUP like), tail to tail inversion (t2tINV), or head to head inversion (h2hINV).

In addition to the aforementioned chromoplexy event, VCaP also harbored a highly complex chromothripsis event on chromosome 5 characterized by clustered SVs with random orientation and alternating copy number changes (Fig. 5C). This chromoplexy event interrupted *TBC1D9B* and *EBF1* at exons, in addition to *IL12B, CPEB4, FAF2, NDUFAF1, MAST4, ADAMTS6, SERF1A, ANKRD31, CNOT6, PDE8B, DMGDH, FAM81B, BTNL9, TMEM232* at introns (Tables S6; Table S7). PC3 also showed evidence of chromothripsis, occurring on multiple chromosomes. In addition to two striking chromothripsis rearrangement clusters on chromosome 5q and 8q, we noted what appeared to be an unusual interchromosomal chromothripsis event bridging chromosomes 1 and 10 (Fig. 5D). This cluster of rearrangements had many of the characteristics of chromothripsis, including numerous rearrangement breakpoints in random orientation with alternative copy number alterations associated with them, except that they were occurring across two separate chromosomes. This was suggestive of an inter-chromosomal rearrangement between chromosomes 1 and 10 in the PC3 cells, with subsequent catastrophic chromothripsis of the newly formed fused chromosomes. All of the chromothripsis breakpoints were chained together, consistent with the observation that SVs in one chromothripsis event only involve one chromosome at a time, with the chromosome in this case being a previously rearranged and fused chromosome 1 and 10 chimera.

Notably, both PC3 and VCaP had hotspot or inactivating *TP53* mutations (Fig S1D); the co-occurrence of *TP53* mutations and the presence of chromothripsis rearrangements in these cell lines (Fig. 3A; Fig. 4C; Fig. 5C,D; Fig. S1D) is consistent with the recent observation from Quigley et al. (10) showing a strong association between *TP53* mutations and presence of chromothripsis events in advanced human PCas. Other cell lines also exhibited evidence of associations between somatic mutations and the SV landscape that have been observed in human PCas. For example, the parental LAPC4 cells harbored a mono-allelic frameshift mutation in *CDK12* and already exhibited a significant number of tandem duplications. The LAPC4-CR cells developed a mutation in the other copy in addition to the mutation in the parental cells, and exhibited even more tandem duplications than parental line, particularly evident on chromosomes 7 and 8, including around the MYC locus (Fig. 6C,D; Fig. S2B, Fig. S4E,F). This tandem duplication molecular phenotype has been previously reported to strongly associate with human PCas harboring *CDK12* mutations (9, 10, 11). Taken together, these associations demonstrate that the cell lines recapitulate many of the associations that have been observed between somatic mutations and structural variants in human PCa.

**Figure 6:**
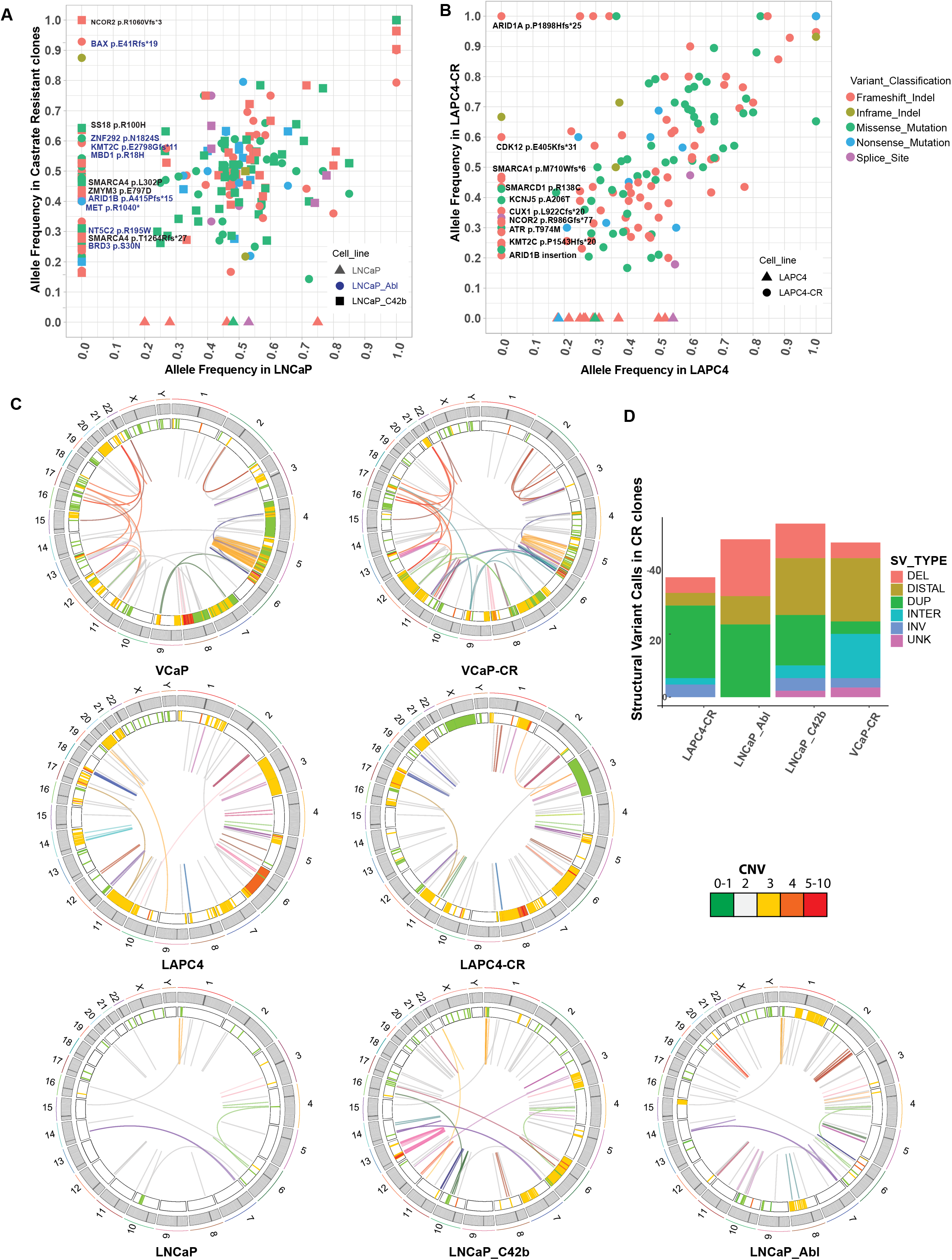
Comparison of LNCaP, LAPC4, and VCaP parental cell lines with their respective CR derivatives. **(A**,**B)** Allele frequencies for each mutation in CR clones and its associated parental lines. Mutations seen along the y-axis represent new mutations arising in the CR lines not found in the parental cells. **(C)** Juxtaposition of SVs in parental and CR clones. Heatmap track beneath chromosome band represents CNV with colors representing copy number. Innermost link track represents large SVs as defined by barcode overlap: complex SVs are colored by chain; simple SVs are gray. **(D)** SVs and CNVs present in each CR subclone stratified by SV type. DEL= deletion, DISTAL= distal, DUP= duplication, INTER= interchromosomal, INV=inversion, UNK=unknown.

### Structural variants and copy number variants disrupt many Cosmic-Longtail genes

Structural variants and CNVs also disrupted a few genes in the *Cosmic-Longtail* list. Structural variant breakpoints landed on exons, introns, or within 5kb from 24 genes in the *Cosmic-Longtail* list (Table S7). Notably, the tumor suppressor *FHIT* was involved in multiple unique duplications and deletions in many cell lines. Specifically, in DU145, two homozygous deletions of 42 kb and 30 kb were found in introns (Table S7) and two CNV=1 of 260 kb and 1.26 Mb were found towards the beginning and the end of the gene (Table S8). LNCaP_Abl and C42b both shared a 300 kb duplication spanning from the middle of *FHIT* towards the middle of its neighbor gene *PTPRG* (Table S7). On top of that, C42b also carried an additional 139 kb deletion early and a 397 kb duplication towards the end of the gene (Table S7). In LAPC4, a 50 kb deletion was found in a middle intron of the gene (Table S7). Meanwhile, in LAPC4-CR, two duplications of 113 kb and 772 kb were found in the middle of the gene, accompanied by a translocation with chromosome 1 to result in potentially a *FHIT-ESRRG* fusion gene (Table S7), and a 6.1Mb CNV=1 towards the beginning of *FHIT* (Table S8). These observations were consistent with the knowledge that *FHIT* locus is one the most fragile in epithelial cells and it is one of the earliest and most frequently altered in human cancer (33). Additionally, the continued SV CNV accumulation in CR clones suggests *FHIT* might play a role not only in cancer initiation but also progression.

On a separate note, despite showing very few putative cancer driver mutations (Fig. 3A; Table S4), MSS cell lines PC3 and VCaP accumulated more alterations on putative cancer driver genes through SVs and CNVs. To name a few deletions, a previously documented 840 kb deletion on chromosome 10 resulted in total *PTEN* loss in PC3 (Table S9), a 50 kb deletion on chromosome 19 fused exon 2 of the MAPK regulator and tumor suppressor *CIC* with exon 14th of *TMEM145* (Table S7), and a heterozygous 55 kb deletion on chromosome 6 removed the first 2 exons of the type I human leukocyte antigen *HLA-A* (Table S7). PC3 also carried an LOH 133 kb duplication on chromosome 14 which duplicated the 5th exon of the DNA damage repair gene *RAD51B* (Table S7). In VCaP, beside the known *TMPRSS2-ERG* fusion, an inverse translocation on chromosome 5 fused the first 3 exons of the oncogenic transcription factor *EBF1* to the later half of the pseudogene ADGRV1, potentially introducing a fusion gene (TableS7). Beside the known 460 kb deletion on chromosome 8 causing total deletion of the tumor suppressor phosphatase *PPP2R2A* in VCaP, a full 620kb deletion on chromosome 4 removed a series of genes involved in chemokine signaling and inflammatory pathways *CXCL3, CXCL2, MTHFD2L, AREG* (Table S9).

### Somatic mutations and SVs associated with resistance to androgen deprivation in PCa cell lines

We next compared the somatic mutational and structural variant landscape in the parental LNCaP, LAPC4, and VCaP cell lines versus their previously established castrate resistant subclones (23, 24, 25) (Fig. 6). While the CR cell lines largely retained the mutations present in their parental cell line confirming that they are truly subclones of those parental lines, they gained many additional mutations not present in the parental cell lines (Fig. 6A,B). Multiple CR clones from MSI parental cell lines gained alterations in genes involved in mediating epigenetic processes. LNCaP_Abl, LNCaP_C42b, and LAPC4-CR all gained mutations on members of the SWI/SNF chromatin remodeling complex. LNCaP_Abl gained a frameshift mutation on *ARID1B*; LAPC4-CR gained a frameshift mutation as a second hit on top of the existing *ARID1A* mutant allele already present on the parental LAPC4 cell line; and LNCaP_C42b gained missense and frameshift mutations on *SMARCA4* and *SS18* respectively. Given how commonly members of the SWI/SNF complex gained mutations in the castrate resistant sublines, we investigated whether these alterations may be more common in metastatic castration resistant cancer compared to primary or castration sensitive metastatic PCa. We queried the cBioPortal database for mutations in any of the 28 SWI/SNF subunits (38) across 5 PCa genomic studies (6, 8, 39, 40, 41). Eleven percent of all patients (n=160 of 1578 total patients) showed mutations in at least one SWI/SNF subunit (Fig. S5A), with more mutations found in castrate resistant PCa than castrate sensitive PCa (Fig. S5B, C).These preliminary observations suggest SWI/SNF mutations potentially play a role in driving castrate resistant PCa, in addition to a recent study reporting high frequency of SWI/SNF gene alterations in neuroendocrine prostate cancer (42). Additional molecular studies should be done to comprehensively determine if these roles in castration resistance are consistently noted and whether the associations are correlative or causative.

Beside SWI/SNF, other epigenetic machinery proteins that gained mutations in the CR cell lines include the bromodomain protein *BRD3*, a histone deacetylase containing complex member *ZMYM3*, a nucleosome remodeling associated DNA helicase *SMARCAD1*, the methyl-binding domain protein *MBD1*, the histone methyl transferase *KMT2C*, and the nuclear co-repressor protein *NCOR2* (Fig. 6A,B, S4A). Genes from other pathways were also newly mutated in the CR cell lines. LAPC4-CR developed biallelic heterozygous mutations in *KCNJ5*, which encodes an inward-rectifier potassium channel protein. It also gained a mutation in the DNA repair gene *ATR*. For genes involved in apoptosis pathways, LNCaP_Abl lost the pro-apoptotic gene *BAX*, and VCaP-CR gained a frameshift mutation on *BCOR*, which is a *BCL6* corepressor (Fig. S4A). It is interesting to note that none of the newly gained mutations in CR clones were on the *AR* gene itself.

CR clones also accumulated more SVs and copy number variants (CNVs), many of which might have contributed to their CR progression (Fig. S4). LNCaP_Abl and LAPC4-CR had more duplications and deletions, whereas VCaP-CR and C42b gained more complex rearrangements. Most notably, 8q24 amplification was found in both LAPC4-CR and LNCaP_Abl. The LAPC4-CR line, which harbored biallelic mutation of *CDK12*, showed multiple tandem duplications around both the *MYC* gene and its enhancer (Fig. S4E); such an association between *CDK12* mutation and tandem duplications around the MYC gene and its enhancer was recently reported in human PCas (10, 11). Whereas LAPC4-CR gained many tandem duplications around the locus spanning both the *MYC* gene and its enhancers, in LNCaP_Abl there were three copy number gains at 8q24 with one spanning the *MYC* enhancer (Fig. S4D). VCaP-CR has an *AR* enhancer amplification, a well-studied recently discovered SV driving castrate resistant phenotype (Fig. S4B)(11, 43). The *AR* enhancer duplication in VCaP-CR occurred in addition to the *AR* amplification already present in VCaP. Lastly, C42b gained a series of complex SVs on chromosome 10, 11, 12, 13, and 14 (Fig. 5C, Table S7). Most notable was a chromothripsis-like complex rearrangement on chromosome 13 (Fig. S4C). These complex rearrangements involved multiple amplifications and translocations within chromosome 13, dysregulating genes within 13q12.11 to 13q13.2 and 13q31.3. Amongst these, *LATS2* encodes a tumor suppressor protein kinase that functions as a positive regulator of *TP53* and as a corepressor of androgen-responsive gene expression (44) *TNFRSF19* is involved in B-catenin signaling, and apoptosis; and *GPC6* controls cell growth and cell division (45, 46). Future mechanistic studies can examine the role of these mutations in driving resistance to androgen deprivation or PCa progression.

## DISCUSSION

Linked-read sequencing allowed allelic phasing of mutations and structural variants in the commonly used PCa cell lines. This approach combines the high throughput and accuracy of paired end short read sequencing with the power of analyzing high molecular weight DNA. This is accomplished by tagging all fragments from an original HMW DNA molecule with unique DNA barcodes that can be used to assign short read sequence data back to the long DNA molecule from which they were derived.

The phasing information of somatic mutations could be used to derive “Gene level haplotypes”, which distinguished between multiple heterozygous mutations on the same or different copies of each gene. These data allowed for the first time a genome-wide analysis of mono-vs. biallelic driver genes alterations in the PCa cell lines, providing a valuable resource for future studies. For instance, CWR22Rv1 had two heterozygous *BRCA2* mutations on opposite alleles, with frameshift mutation T3033Nfs11* on one and clinically recurrent missense mutation V1810I on the other. Although the clinical significance of V1810I is unknown, this knowledge could inform use of CRW22Rv1 in studies of DNA damage/repair, including investigation of synthetic lethality paradigms, in PCa.

The linked read sequencing data are well-suited to identifying rearrangement breakpoints since the molecular barcodes from the original long molecules provide many chances to identify a rearrangement breakpoint as overlap of barcodes from non-contiguous genomic regions. Furthermore, by developing a simple approach, that we term ChainLink, we could leverage the phasing/haplotype information from the linked read data to phase SV breakpoints and identify chained complex rearrangements, including those consistent with chromoplexy and chromothripsis, more directly than previous approaches. These analyses enabled several important insights into these complex rearrangement events. First, we were able to decipher for the first time the precise genome anatomy of several complex rearrangement events, including the highly recurrent *TMPRSS2-ERG* rearrangement present in VCaP cells (Fig. 5A,B), as well as several chromoplexy and chromothripsis events in numerous cell lines that were previously unrecognized (Fig. 4C; Fig. 5; Fig. 6C; Fig. S3, S4; Table S5; Table S6). Second, by phasing all of the individual breakpoints that constituted these complex rearrangements, we showed that the breakpoints all occurred across a single phased allele, and did not represent rearrangements that were staggered independently in different alleles. This provides further evidence in support of the concept that chromoplexy and chromothripsis rearrangements occur in a single concerted event, rather than through accumulation of multiple independent rearrangements.

These analyses also revealed several important insights regarding the relevance of this set of commonly used PCa cell lines to human PCa. First, the cell lines captured a large fraction of recurrently mutated driver genes in human PCa as have been reported in large scale PCa genome sequencing studies (7). Additionally, integrating the phased mutation and SV information, we saw many associations between mutations and SV patterns that have been described in human cancers including: i) the association of *CDK12* mutations with increased propensity for tandem duplications as seen in the LAPC4/LAPC4-CR cell lines; and ii) the association of *TP53* mutations with presence of chromothripsis (PC3 and VCaP cells) (9, 10). Finally, comparing the CR cell lines to the castration sensitive parental cell lines from which they were derived, we found that several mutations in epigenetic factors and apoptosis pathways among others occurred in the CR lines. It will be interesting to see if these classes of alterations are associated with development of castration resistance in men with PCa.

One limitation of this study is that the approaches used for phasing variants and rearrangement breakpoints would not have accounted for aneuploidy. As a result of this, we would be limited in determining the phasing of variants that arose after a specific allele gained a copy. Nonetheless, since we had such a high rate of phasing of the identified somatic mutations, it is likely that most of these mutations arose prior to any copy number gains of those segments.

Another issue to consider is that of cell line drift, which may lead to some differences between the genetic features measured in this study compared to those that may be used in individual labs. We therefore carried out significant validation of these cell line identities through STR genotyping and comparing to the source STR typing information, as well as through careful comparison to mutations and genetic alterations reported in the literature for each cell line. We also report the specific sources of the cell lines used in this study on table S1.

In conclusion, this study comprehensively characterizes allelically phased genomic alterations in the commonly used PCa cell lines, provides important insights into the pathogenesis of complex chromoplexy and chromothripsis events in PCa, and provides a useful resource for future cancer research.

## SUPPLEMENTARY DATA

Supplementary Data are attached.

## ACKNOWLEDGEMENTS

We would like to thank Jennifer Meyer from the SKCCC Experimental and Computational Genomics Core and Lisa Haley from the Pathology Molecular Diagnostics Lab for their help with sequencing runs and quality controls of DNA samples.

## SUPPLEMENTARY MATERIALS

### Supplementary Methods

#### Pubmed Query of Prostate Cancer Cell lines

To assess the number of manuscripts citing the use of all of the 20 distinct prostate cancer cell lines reported in references (1,2,3), we carried out the following query in Pubmed on July 2, 2021:

“DU-145”[All Fields] OR “DU145”[All Fields] OR “PC3”[All Fields] OR (“pc 3 cells”[MeSH Terms] OR (“pc 3”[All Fields] AND “cells”[All Fields]) OR “pc 3 cells”[All Fields] OR “pc 3”[All Fields]) OR (“lncap”[All Fields] OR “lncaps”[All Fields]) OR “ARCAP”[All Fields] OR “DUCAP”[All Fields] OR “LAPC-4”[All Fields] OR “LAPC4”[All Fields] OR “MDA-PCA-2A”[All Fields] OR “MDA-PCA-2B”[All Fields] OR “NCI-H660”[All Fields] OR “22RV1”[All Fields] OR “1013L”[All Fields] OR “PC-346C”[All Fields] OR “PC346C”[All Fields] OR “PSK-1”[All Fields] OR “UM-SCP-1”[All Fields] OR “VCAP”[All Fields]

This resulted in identification of 23,029 citations. We also queried each cell line individually, and noted that the top 7 cited cell lines were PC3, LNCaP, DU145, 22RV1, VCaP, LAPC-4, MDA-PCA-2B, which collectively accounted for 22,920 citations which was determined by the following Pubmed query:

“DU-145”[All Fields] OR “DU145”[All Fields] OR “PC3”[All Fields] OR (“pc 3 cells”[MeSH Terms] OR (“pc 3”[All Fields] AND “cells”[All Fields]) OR “pc 3 cells”[All Fields] OR “pc 3”[All Fields]) OR (“lncap”[All Fields] OR “lncaps”[All Fields]) OR “LAPC-4”[All Fields] OR “LAPC4”[All Fields] OR “MDA-PCA-2B”[All Fields] OR “22RV1”[All Fields] OR “VCAP”[All Fields]

## Supplementary Figures

**Figure S1:**
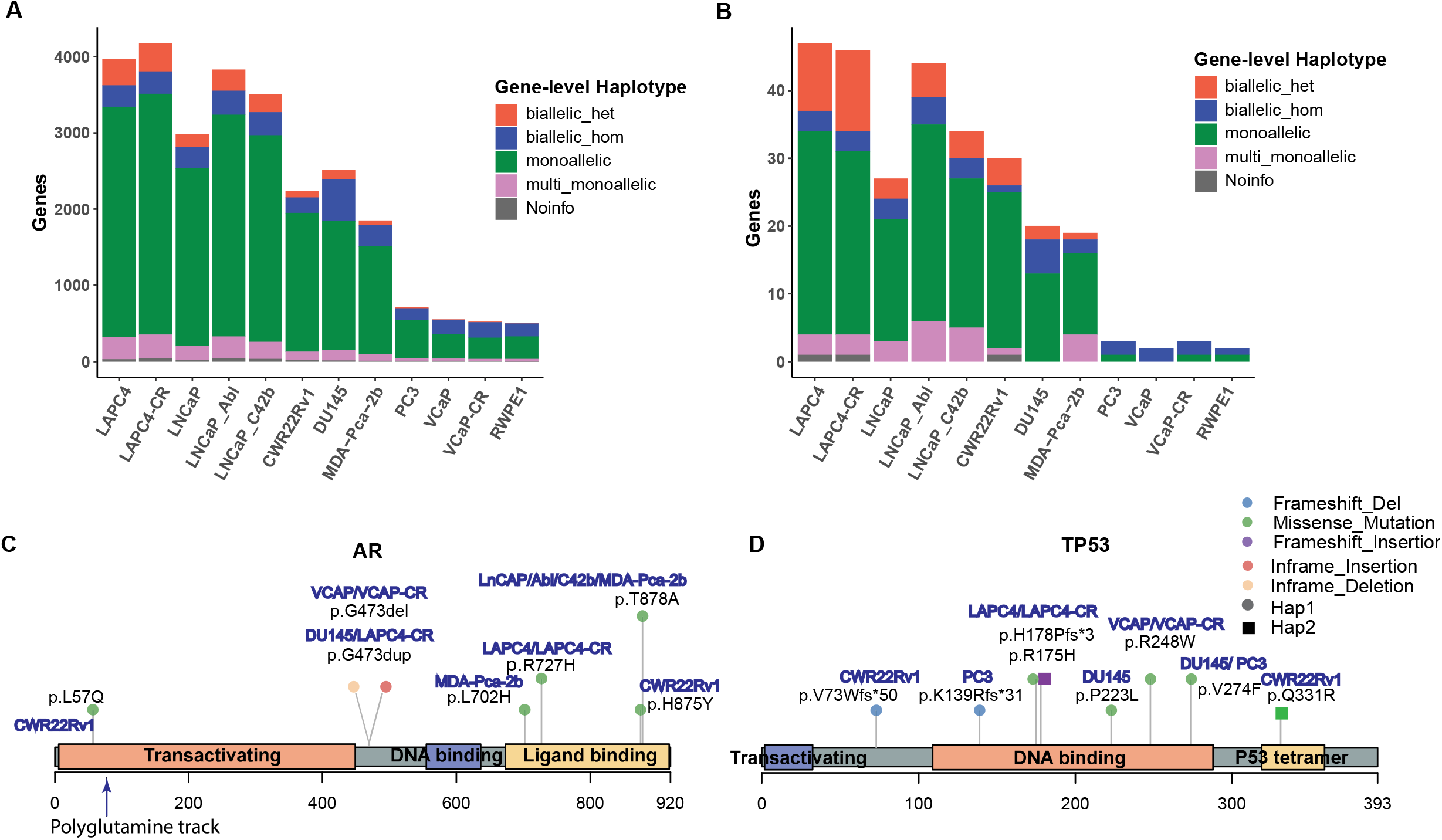
Gene-level Haplotype for putative PCa driver genes. **(A)** Number of somatic mutations across all genes stratified by Gene-level haplotypes. **(B)** Number of somatic mutations across recurrently mutated genes in PCa as compiled in the *Longtail* list, stratified by Gene-level haplotypes. **(C, D)** Lollipop graphs showing Pfam domains and positions of mutations found in AR and TP53 proteins. Shapes and colors of the lollipop head indicate the haplotype and variant classification for each mutation respectively.

**Figure S2:**
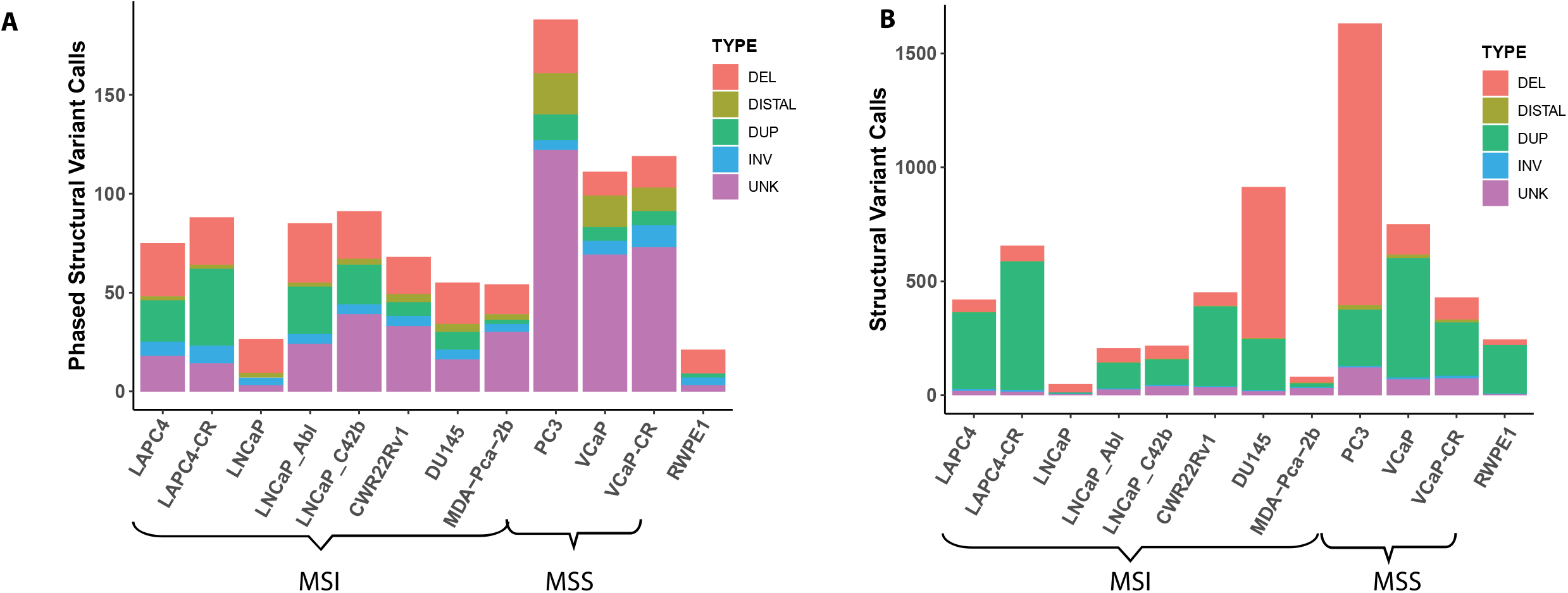
Number of SVs and CNVs present in each cell line. Numbers of phased SVs **(A)** and overall number of SVs and CNVs **(B)** stratified by type. DEL= deletion, DISTAL= distal, DUP= duplication, INV=inversion, UNK=unknown.

**Figure S3:**
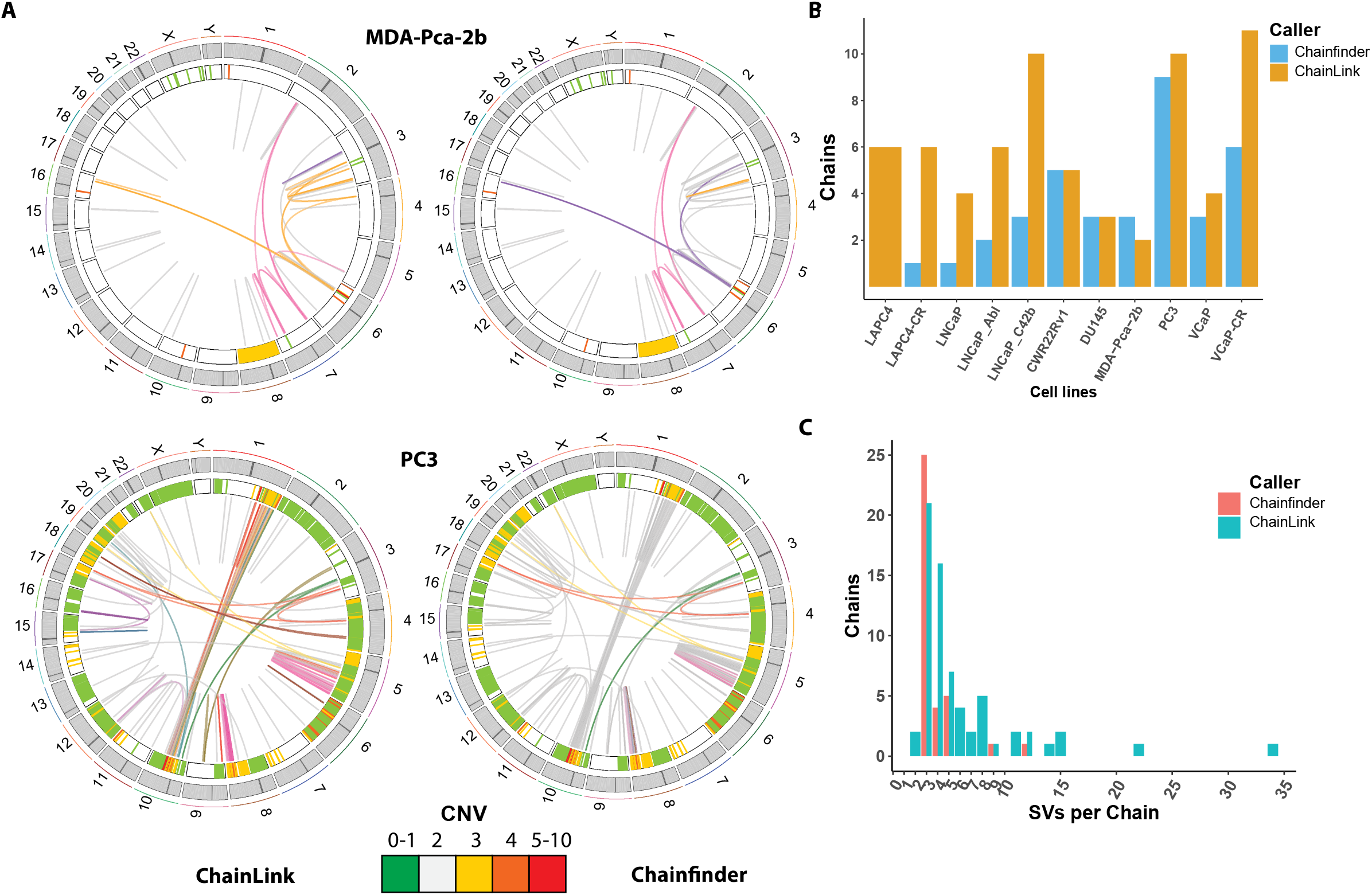
Comparison between ChainLink and ChainFinder in chaining complex SVs in PCa cell lines. **(A)** Complex chained SVs for representative cell lines MDA-PCa-2b and PC3 identified by ChainLink or Chainfinder. Heatmap track beneath chromosome band represents CNV with colors representing copy number. Innermost link track represents large SVs as defined by barcode overlap: complex SVs are colored by chain, with predicted simple SVs identified in gray. **(B)** Number of complex SV chains called by ChainFinder or ChainLink. **(C)** Number of chains and SVs per chain as identified by ChainFinder or Chainlink.

**Figure S4:**
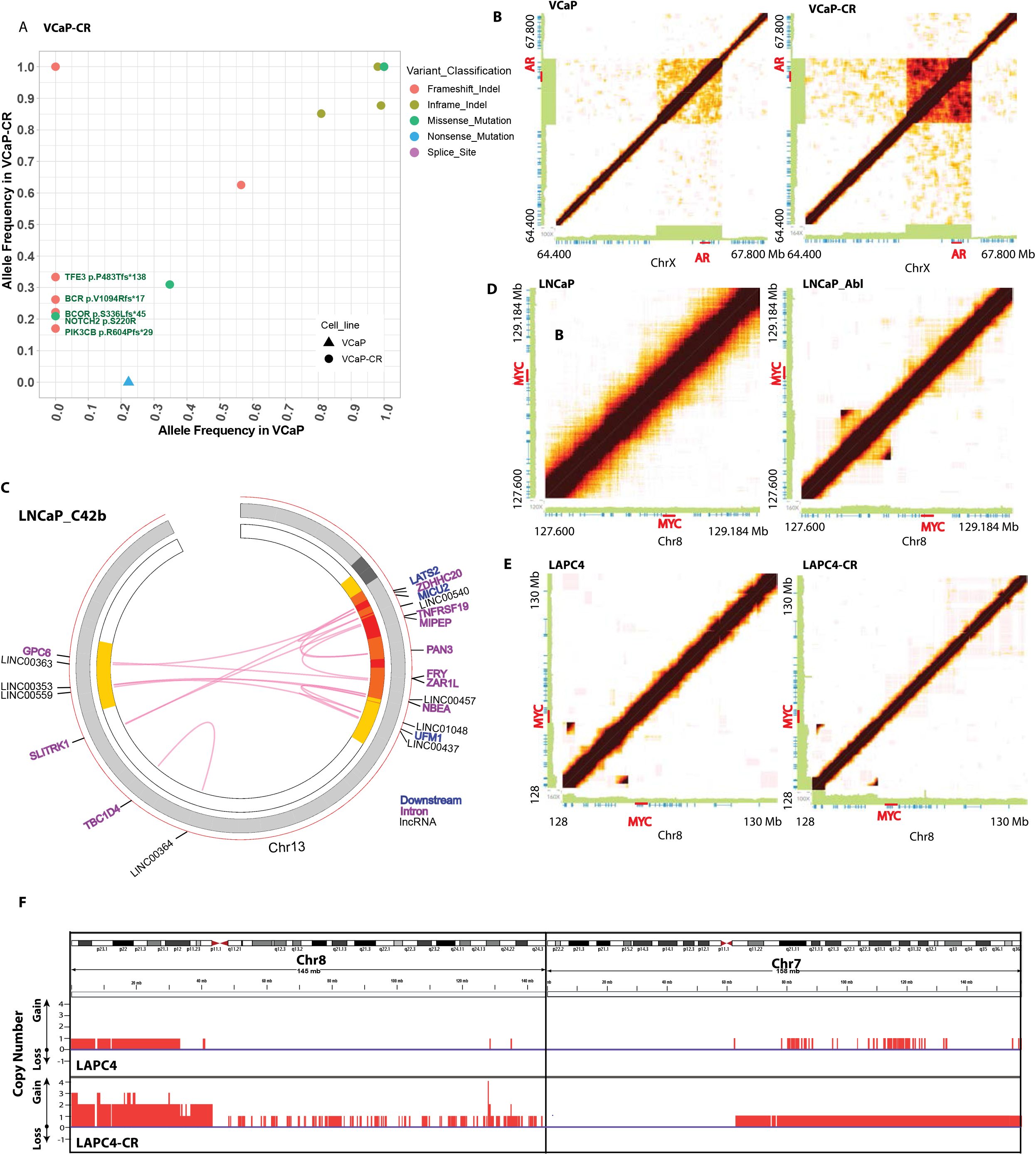
Mutations, structural variants, and copy number variants found in CR clones. **(A)** Allele frequencies for each mutation found in VCaP-CR clones and VCaP. **(B)** VCaP-CR showed copy number gain at *AR* enhancer and *AR* gene compared to VCaP. **(C)** Chromothripsis-like complex SV event on chromosome 13 disrupted many protein-coding introns and lncRNAs in LNCaP_C42b. **(D)** *MYC* enhancer amplification on chromosome 8 in LNCaP_Abl compared to LNCaP. **(E)** Tandem duplications around 8q24 in LAPC4-CR compared to LAPC4. **(F)** Tandem duplication observed in LAPC4 and LAPC4-CR, especially in chromosome 7 and 8. Purple line represents copy number neutral. Red bars above the purple line represent copy number gains.

**Figure S5:**
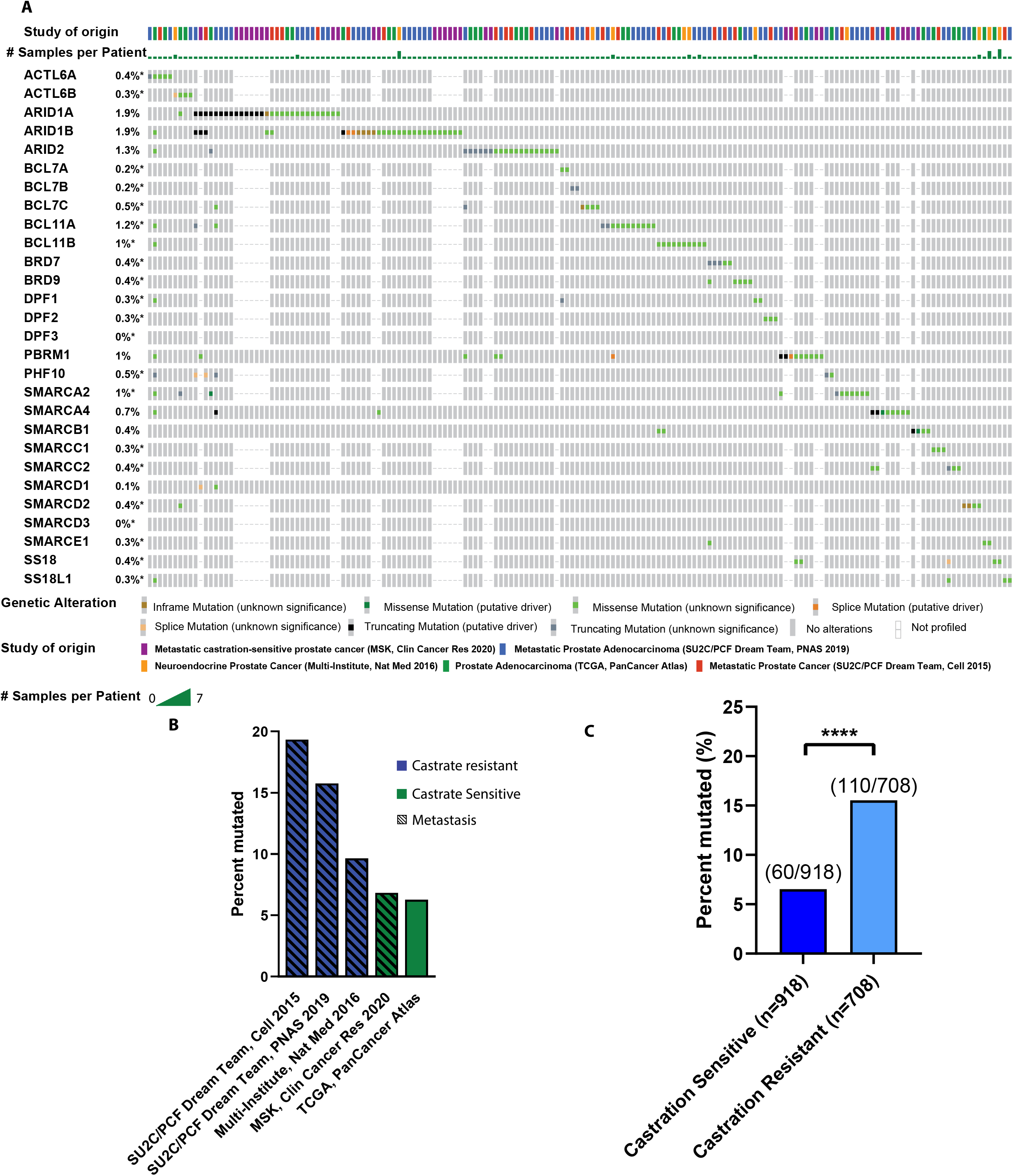
Mutations in SWI/SNF complexes and its association with castrate resistant phenotype. **(A)** Oncoprint showing mutations in SWI/SNF subunits present in 11% of 1578 patients and 1626 samples from 5 Prostate Cancer studies (6, 8, 39, 40, 41). Top color bar represents study origins, colors represent alteration classification. Only samples with at least one SWI/SNF subunit mutated are shown (n=160). **(B)** Frequency of SWI/SNF subunit mutations found in each of the 5 studies queried, from left to right: Metastatic Castration Resistant Prostate Cancer (SU2C/PCF Dream Team, Cell 2015, n=150), Metastatic Castration Resistant Prostate Adenocarcinoma (SU2C/PCF Dream Team, PNAS 2019, n=444), Castrate Resistant Neuroendocrine Prostate Cancer (Multi-Institute, Nat Med 2016, n=114), Metastatic castration-sensitive prostate cancer (MSK, Clin Cancer Res 2020, n=424), Prostate Adenocarcinoma (TCGA, PanCancer Atlas, n=494), total n=1578 patients/1626 samples. **(C)** Percentage of samples with at least one SWI/SNF mutation in castrate resistant (n=708) and castrate sensitive (n=918) PCa patients. SWI/SNF mutations are enriched in Castration resistant PCa compared to Castration sensitive PCa (Chi-squared, p<0.0001).

## Supplementary Tables

**Table S1:** Origins, characteristics, and authentication information of cell lines used in this study

**Table S2:** HMW DNA extraction QC, sequencing depth, and Long Ranger output metrics for each cell line

**Table S3:** Microsatellite instability status, somatic mutation count and somatic mutation rate for each cell line.

**Table S4:** Putative driver mutations in *Cosmic-Longtail* gene list found in our panel of 12 PCa cell lines. Column headings are according to MAF column specification annotated by vcf2maf (18). Frameshift, nonsense, and splice site variants were retained, together with any missense mutations annotated by the Cancer Mutation Census (CMC) list (19).

**Table S6:** List of simple and complex SVs in each cell lines and the phase information used for complex SV calling by ChainLink method

**Table S7:** List of SVs and genes interrupted by SVs in each cell line

**Table S8:** List of CNV in each cell line as called by barcode coverage by Long Ranger

**Table S9:** List of full deletions (CNV=0) in each cell line and associated gene loss

**Table S10**: List of small deletions (> 40bp and < 30kb) detected in each cell line

